# Chromosomal and gonadal sex drive sex differences in lipids and hepatic gene expression in response to hypercholesterolemia and statin treatment

**DOI:** 10.1101/2022.08.29.505758

**Authors:** Carrie B. Wiese, Zoey W. Agle, Peixiang Zhang, Karen Reue

## Abstract

**Background:** Biological sex impacts susceptibility and presentation of cardiovascular disease, which remains the leading cause of death for both sexes. To reduce cardiovascular disease risk, statin drugs are commonly prescribed to reduce circulating cholesterol levels through inhibition of cholesterol synthesis. The effectiveness of statin therapy differs between individuals with a sex bias in the frequency of adverse effects. Limited information is available regarding the mechanisms driving sex-specific responses to hypercholesterolemia or statin treatment.

**Methods:** Four core genotypes mice (XX and XY mice with ovaries and XX and XY mice with testes) on a hypercholesteremic *Apoe*^−/−^ background were fed a chow diet without or with simvastatin for 8 weeks. Plasma lipid levels were quantified and hepatic differential gene expression was evaluated with RNA-sequencing to identify the independent effects of gonadal and chromosomal sex.

**Results:** In a hypercholesterolemic state, gonadal sex influenced the expression levels of more than 3000 genes, and chromosomal sex impacted expression of nearly 1400 genes, which were distributed across all autosomes as well as the sex chromosomes. Gonadal sex uniquely influenced the expression of ER stress response genes, whereas chromosomal and gonadal sex influenced fatty acid metabolism gene expression in hypercholesterolemic mice. Sex-specific effects on gene regulation in response to statin treatment included a compensatory upregulation of cholesterol biosynthetic gene expression on mice with XY chromosome complement, regardless of presence of ovaries or testes.

**Conclusion:** Gonadal and chromosomal sex have independent effects on the hepatic transcriptome to influence different cellular pathways in a hypercholesterolemic environment. Furthermore, chromosomal sex in particular impacted the cellular response to statin treatment. An improved understanding of how gonadal and chromosomal sex influence cellular response to disease conditions and in response to drug treatment is critical to optimize disease management for all individuals.

## INTRODUCTION

Cardiovascular disease (CVD) remains the leading cause of death for both men and women in the United States. CVD mortality for males and females combined declined from 1980 to 2010, but has increased from 2010 to 2019 (the most recent time point for which figures are available) [1]. This is despite extensive knowledge regarding the risk factors for the development of CVDs (including hypercholesterolemia, high blood pressure, and diabetes), and the widespread use of drugs to reduce these risk factors. A fundamental factor that influences CVD susceptibility is biological sex. Men are more susceptible than women to the most common form of cardiovascular disease, coronary artery disease, through age 50 [2,3]. However, women with coronary artery disease before age 50 have a worse prognosis than men, and following menopause, the incidence of disease in women increases and overtakes that in men at advanced ages.

Statin drugs are widely prescribed worldwide to reduce CVD risk by inhibiting hepatic cholesterol synthesis and reducing circulating cholesterol levels. Although statins are largely effective at reducing CVD risk, statin response differs among individuals. In particular, biological females are more likely to experience adverse statin effects, including myopathy and new-onset diabetes [4–6]. Elucidating the mechanisms that underlie sex-specific responses to hypercholesterolemia and statin treatment would be a valuable step toward optimizing CVD prevention and treatment for both sexes.

The liver is the central organ for cholesterol homeostasis and statin action [7]. During the postprandial period, hepatocytes take up lipids from intestinally-derived lipoproteins (chylomicrons) that contain triglycerides and cholesterol esters; during the fasting state, hepatocytes take up fatty acids released from triglyceride stores in adipose tissue. The fatty acids may be oxidized, or fatty acids and cholesterol may be esterified and stored as lipid droplets. Hepatocytes package triglycerides and cholesterol esters into very low-density lipoproteins (VLDL) which are secreted and transport triglycerides and cholesterol to peripheral tissues. The hydrolysis of lipids within the core of liver-derived lipoproteins converts them to intermediate-density lipoproteins, and ultimately, low-density lipoproteins (LDL), which carry most cholesterol in the circulation and are imported into tissues throughout the body via the LDL receptor.

Hepatocytes also synthesize cholesterol and fatty acids de novo. The regulation of hepatic cholesterol biosynthesis is under complex metabolic control by multiple factors, most notably the sterol response element binding protein 2 (SREBP2) transcription factor [8]. When cellular cholesterol levels are low, SREBP2 is activated to induce expression of genes involved in cholesterol synthesis, including that of the rate-limiting enzyme, hydroxymethylglutaryl (HMG)-CoA reductase. When cholesterol accumulates, hepatocytes down-regulate SREBP2 (through the action of the liver X receptor alpha transcription factor) and reduce cholesterol synthesis. The utility of statin drugs in reduction of CVD risk stems from their action as competitive inhibitors of HMG-CoA reductase to reduce endogenous cholesterol biosynthesis [9].

Despite the long history of studies on hypercholesterolemia as a CVD risk factor, and statin drug action, little is known about the mechanistic basis for sex differences in hypercholesterolemia or statin effects. Here, we dissect sex determinants of hepatic transcriptome changes that occur in the hypercholesterolemic state and in response to statin treatment using the Four Core Genotypes (FCG) mouse model. This model independently segregates the development of gonads from the sex chromosomes to generate four sex genotypes: XX and XY mice with ovaries and XX and XY mice with testes [3,10]. We identified specific effects of gonadal sex and chromosomal sex on plasma lipid levels and hepatic gene expression pathways in the hypercholesterolemic state. We further identified the role of biological sex components in statin lipid lowering and regulation of hepatic gene expression. These findings reveal that both gonadal and genetic sex differences contribute to the regulation of a key CVD risk factor and a widely used therapeutic intervention.

## METHODS

### Animals

The *Apoe*^−/−^;FCG mice were maintained at UCLA for >15 generations and were generated as previously described [11,12]. In brief, female C57BL/6J apolipoprotein E knockout (*Apoe*^−/−^) mice (#002052, Jackson Laboratory, Bar Harbor, ME) were crossed with XY^−^ (*Sry*+) *Apoe*^−/−^ male mice to generate XX, XX(*Sry*+), XY^−^, and XY^−^(*Sry*+) offspring on an *Apoe* deficient background. This mouse cohort is referred to as “FCG mice” throughout. At 8–10 wks of age, mice were fed a chow diet containing 5%–6% fat from calories (D1001, Research Diets, New Brunswick, NJ) or the same diet containing pharmaceutical grade simvastatin (0.1 g/Kg; prepared at Research Diets under formulation D11060903i) for 8 wks. The statin concentration was calculated to be the equivalent in mouse of an 80 mg/day dose in human. All mouse studies were conducted in accordance with UCLA Institutional Animal Research Committee (IACUC) approval under a 12-hour light/dark cycle with ad libitum access to food and water. At time of sacrifice, mice were fasted for 5 hrs and tissues were collected and flash-frozen in liquid nitrogen, followed by storage in –80°C.

### Plasma lipid quantitation

Total cholesterol, triglycerides and free fatty acid levels were determined in mouse plasma via enzymatic reactions and colorimetric detection [13].

### RNA-sequencing

RNA from snap-frozen liver samples was isolated with TRIzol (Thermo Fisher Scientific, Waltham, MA). The RNA-seq libraries were generated as previously described [14], including poly (A) RNA selection, RNA fragmentation, oligo(dT) priming with cDNA synthesis, adaptor ligation to double-stranded DNA, strand selection and PCR amplification to produce the final libraries. Index adaptors were used to multiplex samples after quantification (Quibit) and for quality evaluation (4200 TapeStation, Agilent). Sequencing was performed on a NovaSeq 6000 sequencer at the UCLA Technology Center for Genomics & Bioinformatics. Processing of RNA-seq raw data was performed as previously described [14]. Reads were aligned to mouse genome (GRCm38.97) and read counts per gene were generated with STAR. Differential gene expression analysis was performed with DESeq2 (version 1.26, *n* = 3 per FCG genotype and treatment) and a P value < 0.05 demonstrating significance, unless otherwise noted [15]. Pathway analysis was performed for differentially expressed genes with >1.25 fold-change and adjusted p ≤ 0.05 (chow only comparisons) or p ≤ 0.05 (statin treated comparisons) using Enrichr [16–18]. MA plots, PCA plot, and normalized read boxplots were generated in R. RNA-seq data has been deposited in GEO (Accession number GSE202977).

### Quantitative real-time PCR

For gene expression quantification, RNA was reverse transcribed to cDNA with iScript reverse transcriptase (Bio-Rad, Hercules, CA) and quantified by quantitative real-time PCR (RT-PCR) with SsoAdvanced SYBR Green Supermix on Bio-Rad CFX Opux 384 Real-Time PCR Detection System. All real-time PCR data was normalized to *Rplp0* (also known as *36B4*) gene expression. Supplemental Table 1 contains all primer sequences used in this study.

### Statistical Analyses

Graphpad Prism 9 and RStudio were used for all statistical analyses and data representation. FCG gene expression was analyzed by two-way ANOVA with main factors of gonads (testes vs. ovaries) and sex chromosome complement (XX vs. XY), and assessment of an interaction between the two factors.

## RESULTS

### Effects of gonadal sex on lipid levels are abolished following statin treatment

To assess the effects of sex components on hypercholesterolemia, we made use of the apolipoprotein E (apoE)-deficient (*Apoe*^−/−^) mouse model, which is widely used in the study of hypercholesterolemia and atherosclerosis. *Apoe*^−/−^ mice typically develop hypercholesterolemia (300–600 mg/dL) without dietary intervention, which is associated with elevated levels of LDL and VLDL, as well as reduced high-density lipoproteins (HDL) (reviewed in [19]). We generated *Apoe*^−/−^ FCG mice on a C57BL/6 background (see Methods) to evaluate the role of gonadal and chromosomal sex in response to hypercholesterolemia and statin treatment (Fig. 1A, left panel). Comparison of the two genotypes with XX chromosomes (XX with ovaries and XX with testes) vs. those with XY chromosomes (XY with ovaries and XY with testes) allows for detection of sex chromosome effects (Fig. 1A, middle panel). Comparison of mice with ovaries (XX or XY with ovaries) vs. those with testes (XX or XY with testes) allows for detection of gonadal effects. The analyses that we performed include plasma lipid levels and assessment of the liver transcriptome by RNA-seq and differential gene expression with pathway analyses (Fig. 1A, right panel).

**Figure 1.**
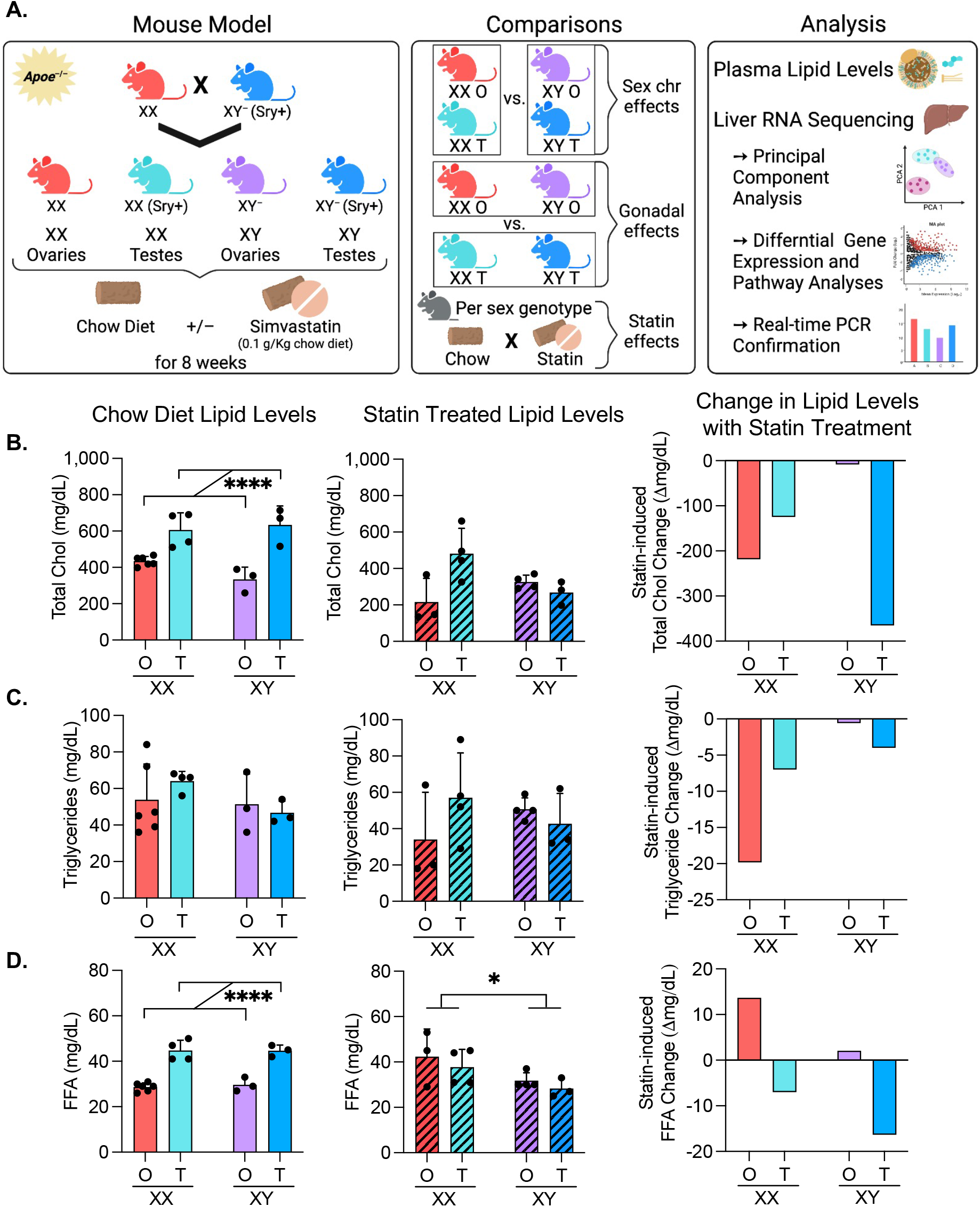
Study design using hypercholesterolemic Four Core Genotypes mice, and sex determinants of plasma lipid level response to statin. (**A**) Experimental design to assess the contribution of sex components to lipid levels and hepatic transcriptome. *Left*, generation of FCG mouse cohorts on hypercholesterolemic (*Apoe*^−/−^) genetic background. *Center*, group comparisons made throughout the study. *Right*, a summary of analyses performed. (**B**) Total cholesterol levels, (**C**) triglyceride levels, (**D**) and free fatty acid levels in plasma of *Apoe*^−/−^ FCG mice on chow diet (left panel), treated with simvastatin (center panel), and the statin-induced change in lipid levels (right panel). For the lipid levels, values represent mean ± standard deviation and were analyzed by two-way ANOVA for gonad and sex chromosome effects indicated by brackets. Denoted significance values: *P<0.05, **** P<0.0001.

*Apoe*^−/−^ FCG mice were fed chow diet with or without simvastatin for 8 wks. On chow diet, all genotypes were hypercholesterolemic due to apo E deficiency, but mice with testes had higher cholesterol levels than those with ovaries, regardless of sex chromosome type (Fig. 1B, left). After statin treatment, there were no significant sex differences in the absolute cholesterol levels (Fig. 1B, middle). However, the sex genotype influenced the statin-induced change in cholesterol levels. Within-genotype reductions in cholesterol levels occurred only in mice having the standard sex genotypes of XX with ovaries or XY with testes; XX mice with testes and XY mice with ovaries did not exhibit a significant drop in cholesterol levels with statin treatment (Fig. 1B, right). In contrast to cholesterol levels, triglyceride levels did not vary across sex genotypes on chow or after statin treatment (Fig. 1C).

Free fatty acid levels in hypercholesterolemic mice were influenced by gonadal type, with the presence of testes associated with higher levels (Fig. 1D, left). After statin treatment, free fatty acid levels were influenced by chromosomal sex, with XX mice having higher levels than XY mice (Fig. 1D, middle). As with cholesterol levels, statin-induced changes in free fatty acid levels occurred only in the standard sex genotypes (XX mice with ovaries and XY mice with testes) (Fig. 1D). However, while statin reduced cholesterol levels in both of these genotypes, fatty acid levels were increased by statin in XX mice with ovaries and reduced in XY mice with testes. Overall, statin altered plasma cholesterol and free fatty acid levels in a sex-dependent manner with the most robust effects in mice having XX chromosomes paired with ovaries or XY chromosomes paired with testes.

### Gonadal and chromosomal sex independently impact the hepatic transcriptome in hypercholesterolemic mice

To identify genes and pathways that influence sex differences in hypercholesterolemia and statin response, we performed RNA-sequencing of liver from *Apoe*^−/−^ FCG mice. To visualize the impact of gonads, sex chromosomes, and statin treatment on gene expression, we performed principal component analysis (PCA) of the RNA-seq data (Fig. 2A). The PCA generated separate clusters for gonadal type based on principal component 1, which accounted for 58% of the variation in the dataset. Within each gonadal type, the samples were separated into individual clusters for XX and XY sex chromosome complement based on principal component 2, which accounted for 23% of the variation within the dataset. Chow and statin treatments clustered together within each genotype indicating that statin treatment had a smaller impact on gene expression than sex genotype. We performed differential expression analysis with DESeq2 [20]; boxplots of the transformed counts show similar total counts across all 24 samples in the analysis (Fig. 2B).

**Figure 2.**
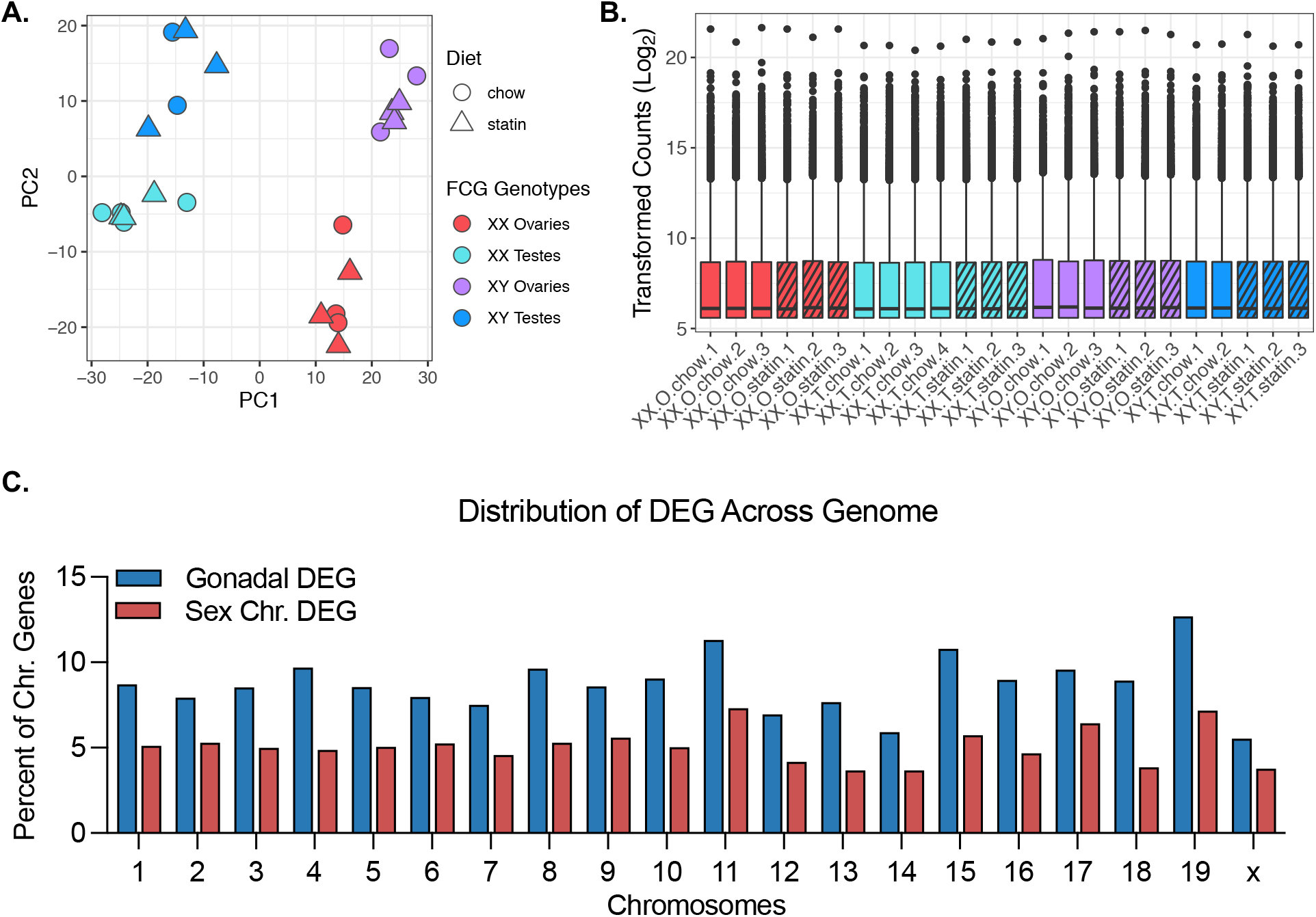
Characterization of *Apoe*^−/−^ FCG liver RNA-seq dataset. (**A**) Principal component analysis of liver RNA-seq data from *Apoe*^−/−^ FCG mice fed chow diet without or with simvastatin for 8 wks. (**B**) Boxplots representing transformed RNA-seq counts across all 24 liver samples. (**C**) The genome-wide distribution of genes found to be differentially regulated by presence of ovaries vs. testes (gonadal DEG), or by presence of XX vs. XY chromosomes (Sex Chr DEG). DEG, differentially expressed genes.

We assessed the influence of sex genotypes on gene expression in the hypercholesterolemic state without statin treatment. A genome-wide analysis of differential gene expression (≥1.25-fold difference, adj p<0.05) identified 3223 genes with differential expression in mice with ovaries compared to testes, and 1390 genes with differential expression in XX compared to XY mice (Supplemental Table 2). We found that 5.9–12.7% of genes on each autosome and 5.5% of genes on the X chromosome had different expression levels in mice with ovaries compared to mice with testes (Fig. 2C, blue bars). The XX vs. XY chromosome complement conferred differential expression of 3.7–7.3% of genes/autosome, as well as 3.8% of genes on the X chromosome (Fig. 2C, red bars). The majority of Y chromosome genes are expressed at very low levels in liver and were therefore not included in this analysis. This analysis reveals that gonadal and chromosomal sex each influence expression of genes that map across all autosomes as well as on the X chromosome.

We determined the functional classification of genes that were differentially expressed based on gonadal or chromosomal sex in the liver of hypercholesterolemic mice by pathway enrichment analysis [16–18]. The 1972 DEG with elevated expression in mice with ovaries compared to testes were enriched for genes in the immune system, cell cycle, and cell signaling pathways of the 49 significant pathways identified (Fig. 3A,B, Supplemental Table 2-3). The 1251 DEG elevated in the presence of testes were enriched in functions that include the unfolded protein response and endoplasmic reticulum (ER) stress pathways (Fig. 3A,B, Supplemental Table 2-3). These results demonstrate that distinct hepatic cellular processes are influenced at the gene expression level by presence of ovaries or testes.

**Figure 3.**
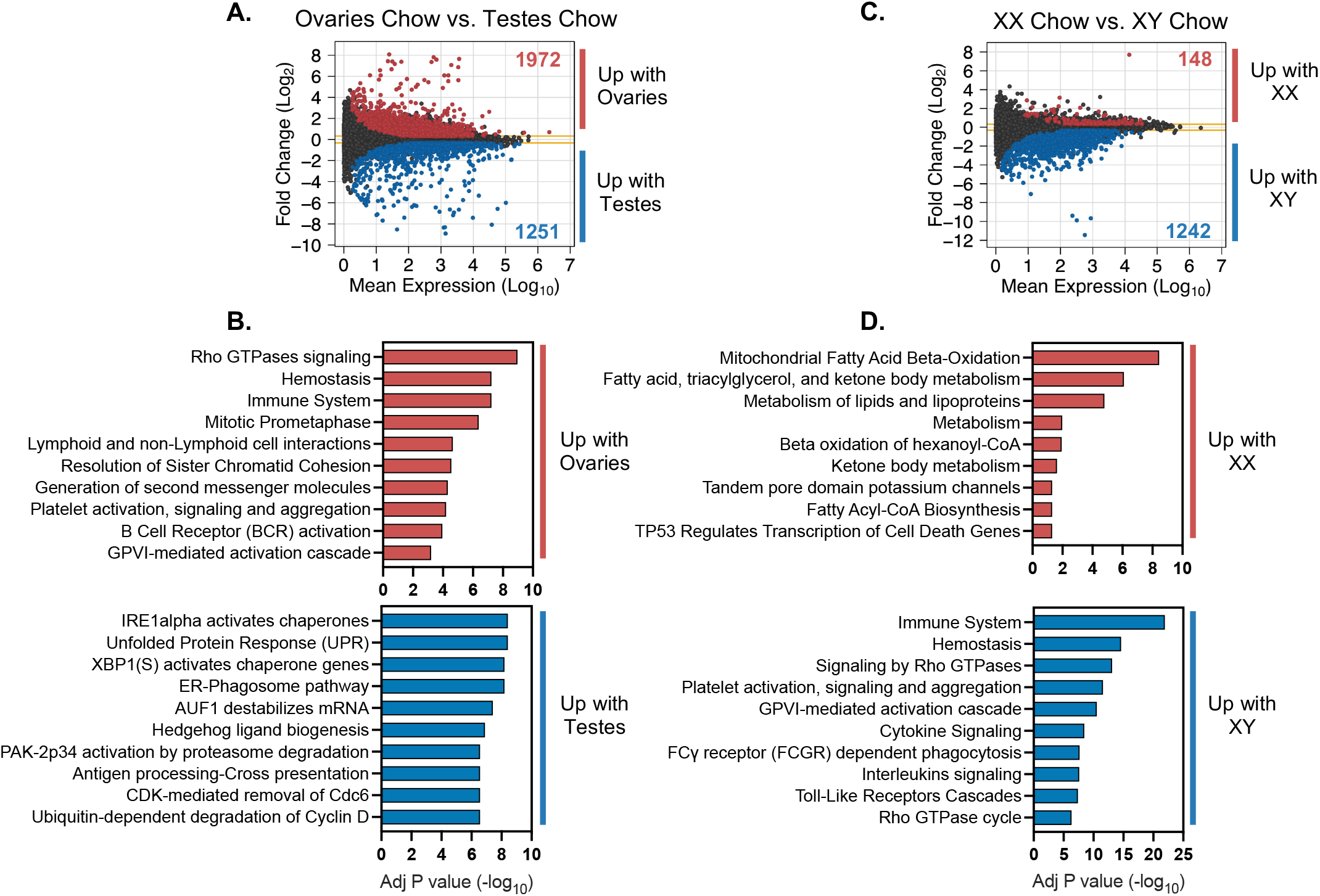
Gonadal and chromosomal sex impact gene expression in *Apoe*^−/−^ FCG liver. MA plot shows mean expression and fold change of genes with differential expression in (**A**) mice with ovaries compared to mice with testes (adjusted P<0.05, fold-change >1.25) and (**B**) the top 10 significant cellular pathways enriched in the differentially expressed genes (adjusted P<0.05). MA plot shows mean expression and fold change of genes with differential expression in (**C**) XX mice compared to XY mice (adjusted P<0.05, fold-change >1.25) and (**D**) the top 10 significant cellular pathways enriched in the differentially expressed genes (adjusted P<0.05).

We also assessed the functional enrichment of genes with differential expression in mice with XX (with ovaries or testes) vs. XY chromosomes (with ovaries or testes). The majority of DEG due to sex chromosome complement (1242 of 1390 genes) had elevated expression in XY compared to XX mice (Fig. 3C, Supplemental Table 2). The XY DEG were enriched in immune system, Rho GTPase signaling, and other cell signaling pathways (Fig. 3D, Supplemental Table 3). Only 148 genes were expressed at higher levels in XX compared to XY liver, but these were enriched for genes associated with fatty acid oxidation in mitochondria and other aspects of fatty acid metabolism (Fig. 3D, Supplemental Table 3).

### ER stress response gene expression is elevated in mice with testes

Since ER stress-related pathways were the predominant pathways upregulated in mice with testes, we further evaluated the regulation of specific genes within these pathways. The ER plays key roles in protein synthesis and trafficking as well as the synthesis of lipids, including sterols, phospholipids, and triglycerides. Disruption of ER homeostasis induces stress responses that are associated with the development and progression of metabolic diseases including obesity, fatty liver, type 2 diabetes and atherosclerosis [21,22]. Of the 92 protein encoding genes in the unfolded protein response pathway (Reactome R-HSA-381119) [23], 36 genes were differentially expressed in mice with testes vs. ovaries, and 7 genes were influenced by sex chromosome complement (two-way ANOVA, p<0.05) (Fig. 4A). We confirmed the expression patterns of representative genes via real-time PCR in a greater number of mouse liver samples. Consistent with RNA-seq data, the expression levels of *Psmd14*, *Hyou1*, *Dnjab11*, and *Sec61a1* were upregulated in the livers of mice with testes compared to mice with ovaries (Fig. 4B). These data suggest that gonadal sex is a key determinant of sex differences in gene expression of ER stress-associated pathways in hypercholesterolemic liver, while chromosomal sex has only a minor impact.

**Figure 4.**
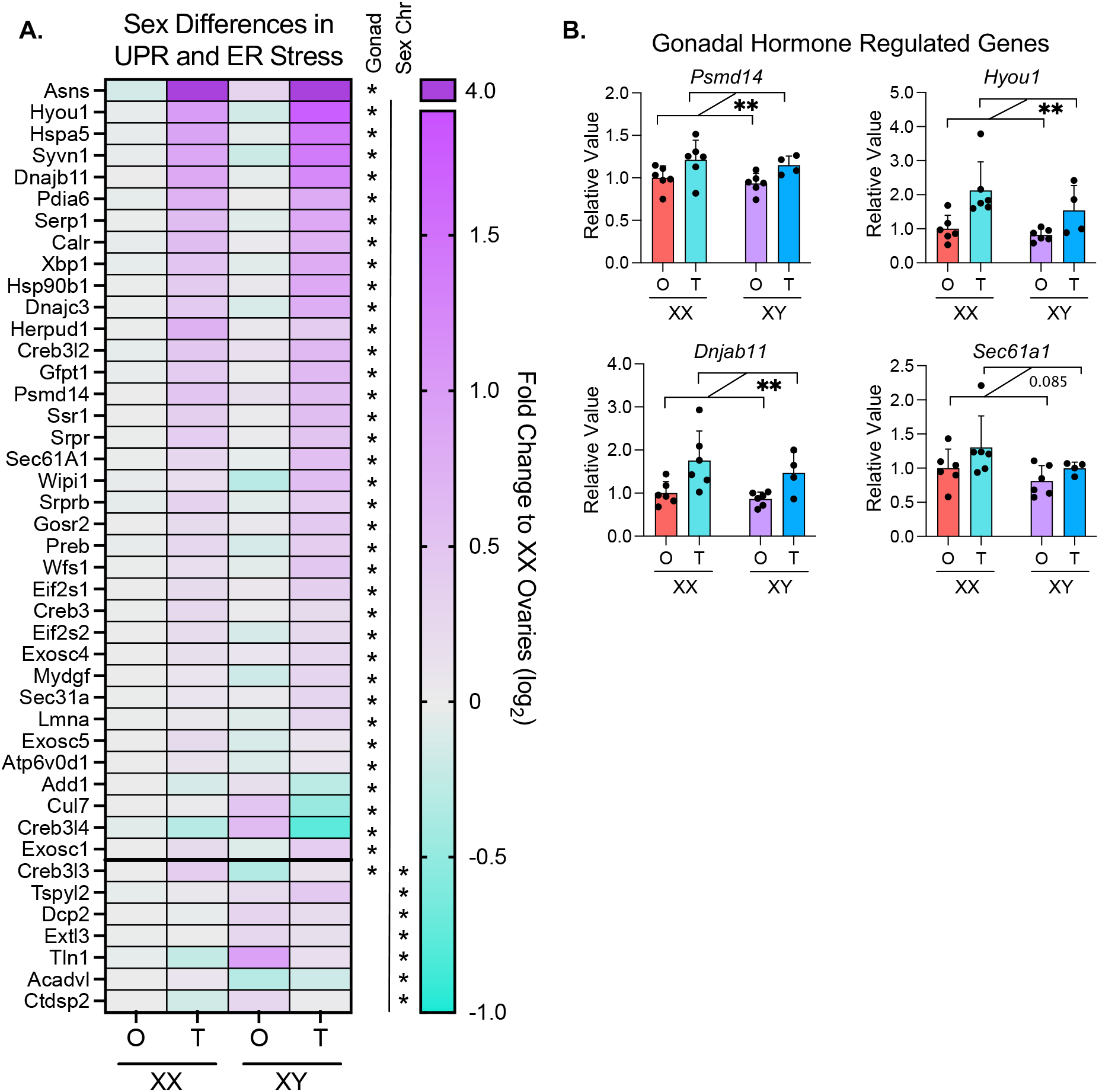
Genes in ER stress and UPR pathways show increased expression in mice with testes. (**A**) Heatmap displays relative hepatic expression levels of genes related to ER stress and unfolded protein response in *Apoe*^−/−^ FCG genotypes. Expression of each genotype was relative to that in XX mice with ovaries. *P<0.05 for gonadal or chromosomal sex by two-way ANOVA. (**B**) Real-time PCR quantification of representative ER stress genes from (A) (*Psmd14*, *Hyou1*, *Dnjab11*, and *Sec61a1*) in liver of *Apoe*^−/−^ FCG mice. **P<0.01 by two-way ANOVA. N=4–6.

### Fatty acid metabolism genes are independently regulated by gonadal and chromosomal sex

Our gene enrichment analyses demonstrated that gonadal and chromosomal sex each impact expression of lipid metabolism genes. In particular, XX vs. XY chromosomes influenced fatty acid metabolism and beta oxidation genes, and testes vs. ovaries influenced peroxisomal lipid metabolism (Fig. 3B,D; Supplemental Table 3). We further investigated the sex component regulation of genes involved in fatty acid synthesis and oxidation across the FCG genotypes. Of the 37 genes in the fatty acid synthesis pathway (Reactome R-HSA-75105.7), 21 genes were regulated by gonadal sex, sex chromosome complement, or a combination of both sex factors in livers of hypercholesterolemic mice (Fig. 5A). Gonadal sex influenced expression of 10 fatty acid synthesis genes. *Elovl3* was strongly increased by the presence of testes, but the majority of the other 10 gonadally regulated fatty acid synthesis genes were down-regulated in mice with testes. The sex chromosome complement influenced expression of 7 fatty acid synthesis genes, with the majority upregulated in XY compared to XX mice. Four fatty acid synthesis genes showed more complex regulation, with significant effects of both gonadal and chromosomal sex.

**Figure 5.**
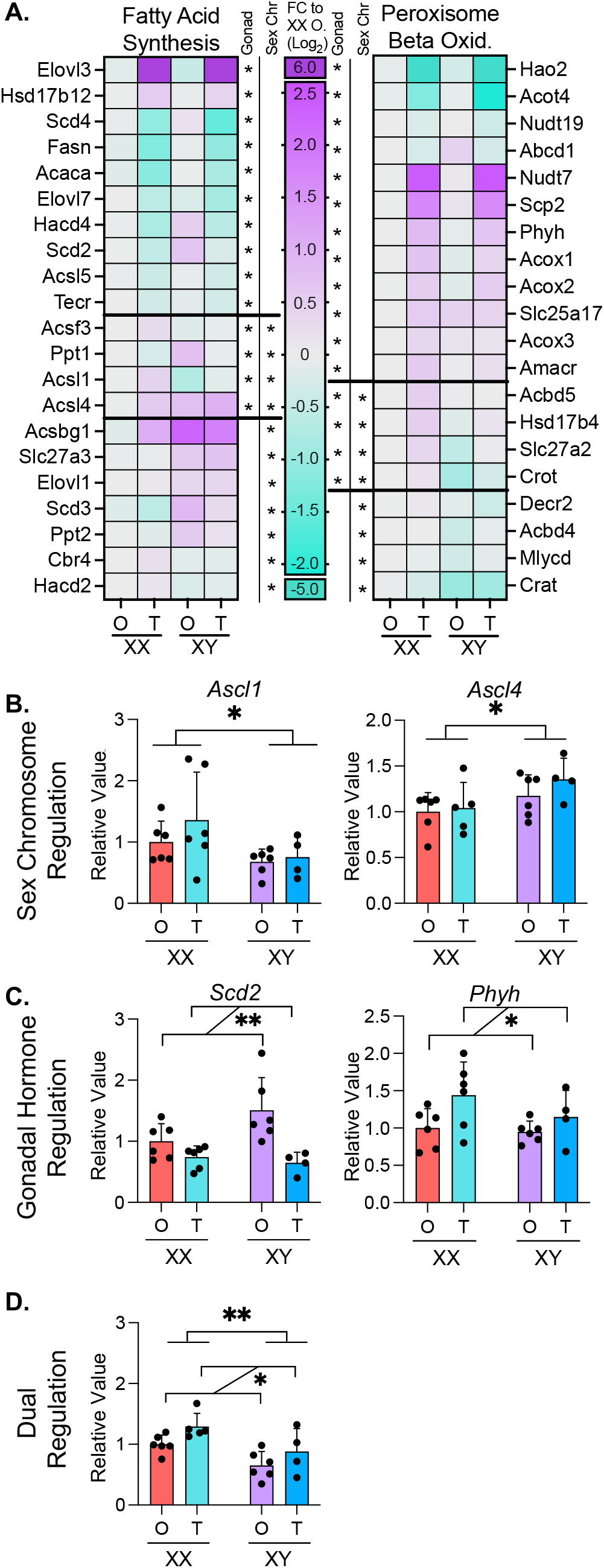
Gonadal and chromosomal sex regulate fatty acid metabolism. (**A**) Heatmaps display relative hepatic expression levels of genes related to fatty acid synthesis and peroxisomal beta oxidation in *Apoe*^−/−^ FCG genotypes. Expression of each genotype was relative to that in XX mice with ovaries. *P<0.05 for gonadal or chromosomal sex by two-way ANOVA. Real-time PCR quantification of representative genes from (A) showing (**B**) chromosomal sex effects (*Acsl1* and *Acsl4*), (**C**) gonadal sex effects (*Scd2* and *Phyh*) and (**D**) both gonadal and chromosomal sex effects (*Slc27a2*). Two-way ANOVA with *P<0.05 and **P<0.01. N=4–6.

Peroxisomal fatty acid oxidation genes were also largely influenced by gonadal sex, but in a pattern distinct from the fatty acid synthesis genes. Of 29 peroxisomal lipid metabolism-related genes (Reactome R-HSA-390918.5), 12 were regulated by gonadal sex, primarily with increased expression due to testes compared to ovaries (Fig. 5A). Four of the 29 peroxisomal oxidation genes were expressed at lower levels in mice with XY compared to XX chromosomes.

We confirmed patterns of representative fatty acid metabolism genes identified by RNA-seq in a larger number of mice by real-time PCR. The acyl-CoA synthetase genes *Acsl1* and *Acsl4* were both regulated by sex chromosome complement, but in opposing directions: XX chromosomes promoted higher expression of *Acsl1*, while XY chromosomes promoted higher expression of *Acsl4*. (Fig. 5B). *Scd2* and *Phyh* were oppositely regulated by gonadal sex, with ovaries promoting higher *Scd2* expression, and testes enhancing *Phyh* expression (Fig. 5C). *Slc27a2* exhibited combined gonadal and sex chromosome regulation with higher expression in XX mice compared to XY mice, but also higher expression in mice with testes compared to mice with ovaries (Fig. 5D). Overall, fatty acid metabolism exhibits complex sex-dependent regulation with fatty acid biosynthesis and peroxisomal degradation impacted in opposite directions by gonadal and chromosomal sex.

### Sex-dependent transcriptional response to statin treatment is influenced by XY sex chromosome complement and testes

To identify sex-dependent effects of statin, *Apoe*^−/−^ FCG mice received simvastatin in chow diet for 8 weeks. The role of sex components on statin-induced alterations in the hepatic transcriptome were identified by comparing statin-treated mice to chow diet controls within each genotype. Statin treatment influenced a similar number of genes in mice with testes (949 DEG between statin and chow) and ovaries (939 DEG between statin and chow) (Fig. 6A). However, only ~5% (94) of the genes altered by statin treatment were shared by mice with ovaries and testes, indicating distinct target genes for statin-associated regulation depending on gonadal type. We analyzed genes with up and down regulation in response to statin (Fig. 6B,C) for functional enrichment. In mice with testes, statin down-regulated genes were enriched for fatty acid metabolism pathways, and statin upregulated genes were enriched for cholesterol biosynthesis pathways (Fig. 6D). In mice with ovaries, statin-regulated genes did not show enrichment for specific biological pathways (Fig. 6E).

**Figure 6.**
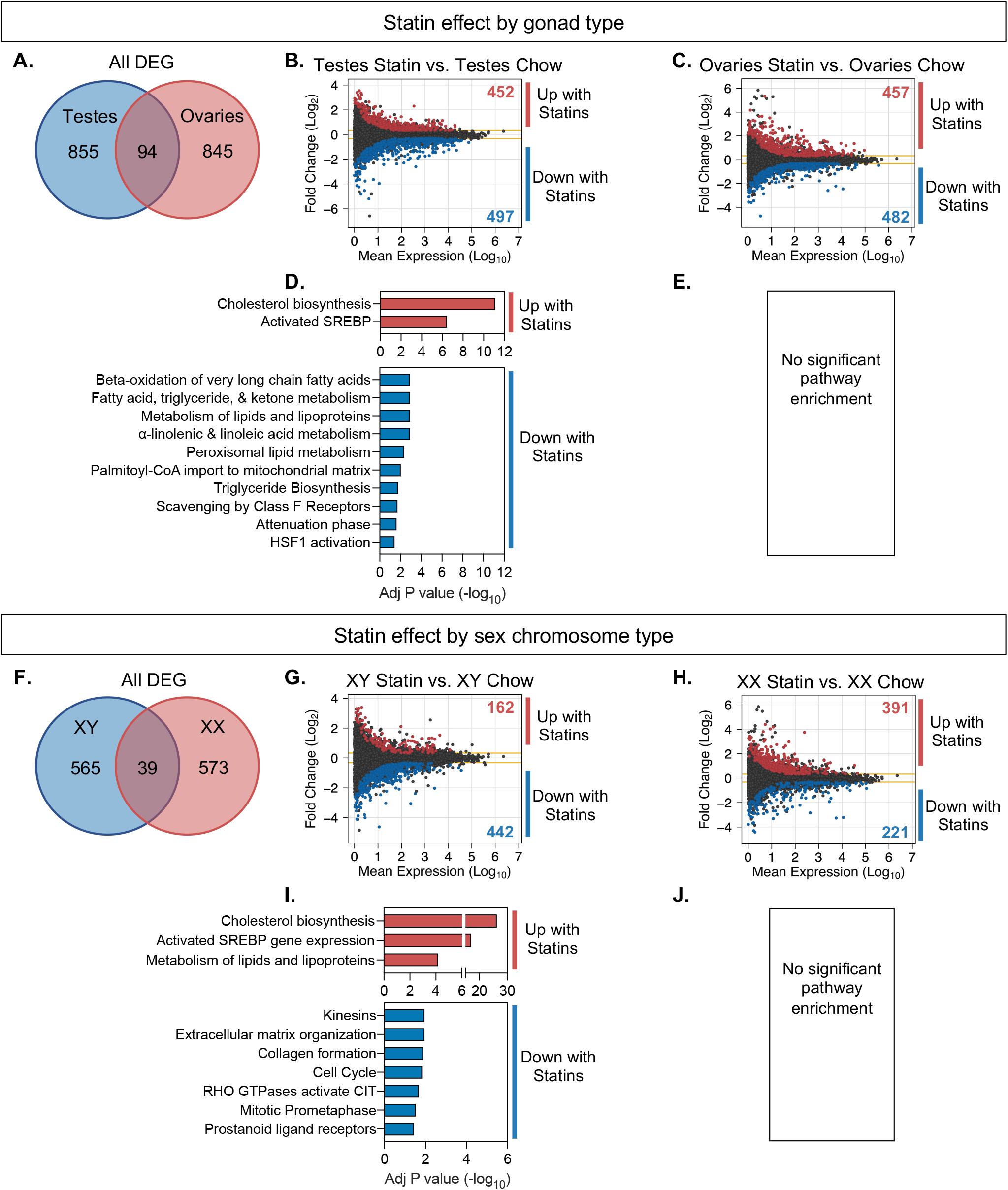
Statin effects on hepatic gene expression are influenced in distinct ways by gonadal and chromosomal sex. (**A**) Venn diagram shows number of genes altered by statin treatment depending on presence of ovaries or testes, and the minimal overlap between the two. MA plots display mean expression levels and fold-change of differentially expressed genes (DESeq2, >1.25-fold altered expression, P<0.05) in (**B**) statin-treated mice with testes compared to chow diet controls and (**C**) statin-treated mice with ovaries compared to chow diet controls. Significantly enriched cellular pathways for statin-induced gene expression changes in (**D**) mice with testes and (**E**) mice with ovaries (no enriched pathways). (**F**) Venn diagram shows number of genes altered by statin treatment depending on presence of XX or XY chromosomes, and the minimal overlap between the two. MA plots display mean expression levels and fold-change of differentially expressed genes (DESeq2) in (**G**) statin-treated XY mice compared to chow diet controls and (**H**) statin-treated XX mice compared to chow diet controls. Significantly enriched cellular pathways for statin-induced gene expression changes in (**I**) XY mice and (**J**) XX mice (no enriched pathways).

Statin-induced gene regulation was also influenced by chromosomal sex. Compared to mice fed a chow diet, statin treatment altered expression of 605 genes in XY mice and 612 genes in XX mice, but only 39 (~6%) of the dysregulated genes were shared between the two sex chromosome complement types (Fig. 6F). In mice with XY sex chromosomes, statin treatment upregulated 162 genes and down-regulated 442 genes. In mice with XX sex chromosomes, statin treatment upregulated 391 genes and down regulated 221 genes (Fig. 6G,H). For XY mice, statin upregulated genes were enriched for cholesterol biosynthesis pathways, while XX mice had no significant pathway enrichment for the statin-induced DEG (Fig. 6I,J).

### Statin treatment induces cholesterol biosynthesis gene expression only in mice with XY sex chromosomes or testes

Statin drugs bind to the rate-limiting enzyme in cholesterol biosynthesis (HMG CoA reductase) to inhibit its activity. The complex feedback mechanisms that control HMG CoA reductase levels can lead to enhanced transcription of the corresponding gene, as well as other genes within the pathway, in response to statin [24,25]. The subsequent upregulation of HMG CoA reductase protein levels may reduce the effectiveness of statin drugs. Our data above suggested that statin-induced compensatory upregulation of cholesterol synthetic gene expression is driven by testes and XY chromosome complement. We investigated further by assessing the effect of gonadal and chromosomal sex on expression of genes in the cholesterol biosynthetic pathway in response to statin. Statin upregulated 18 of 25 cholesterol biosynthetic genes in the liver of XY mice, with little or no increase in XX liver (Fig. 7A). The increased expression in XY mice was most pronounced in XY mice with testes, suggesting an additional impact from gonadal sex.

**Figure 7.**
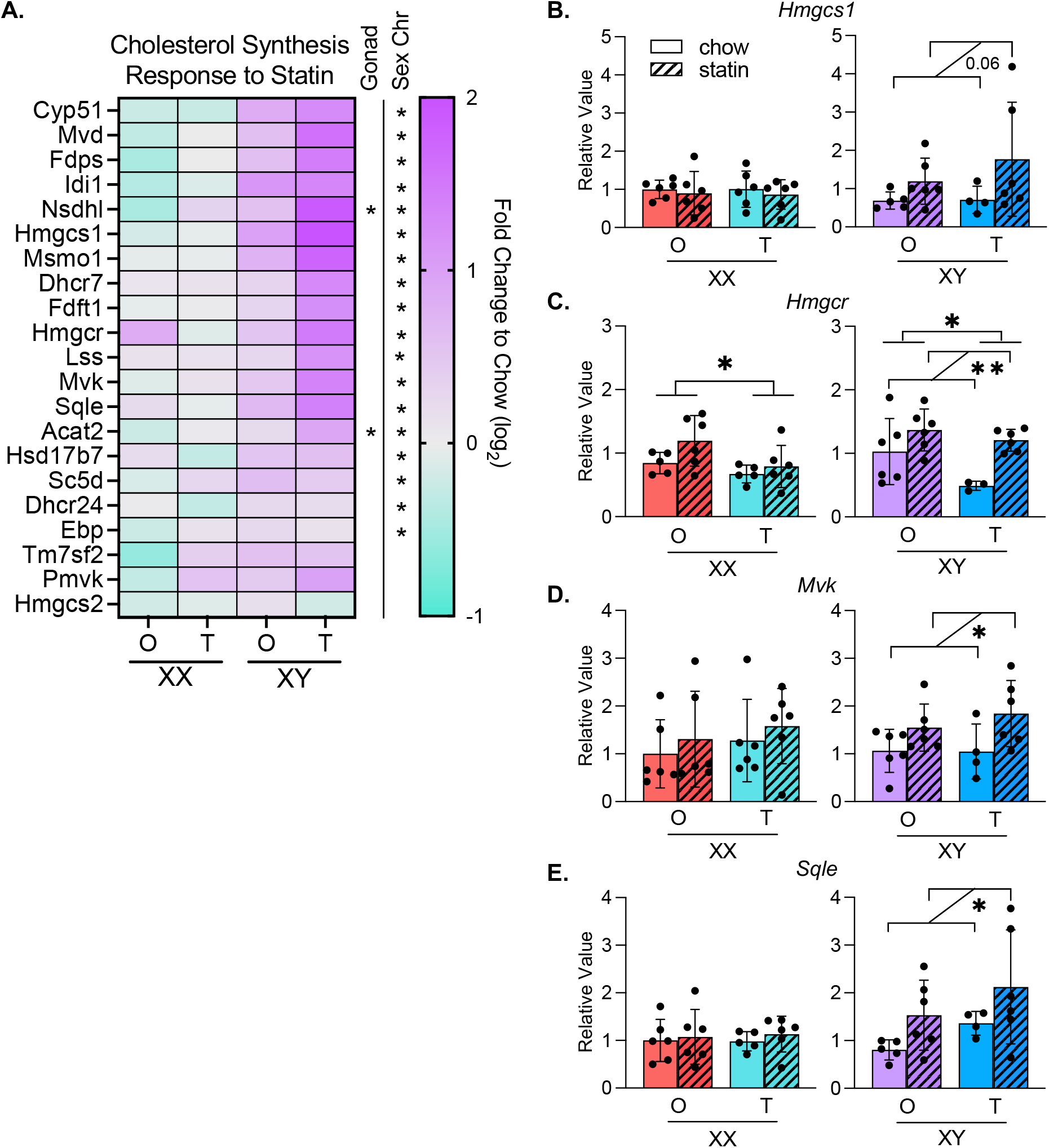
Cholesterol biosynthesis pathway gene expression is up-regulated in XY mice in response to statin. (**A**) Heatmap displays relative hepatic expression levels of genes altered by sex within the cholesterol biosynthesis pathway as fold-change of statin treated compared to chow diet for each of the Four Core genotypes. *P<0.05 by two-way ANOVA. **(B–E)** Real-time PCR quantification of representative genes from (A) illustrates the effects of gonadal and/or chromosomal sex on statin-induced alterations in gene expression in the cholesterol biosynthesis pathway. *P<0.05 and **P<0.01 by two-way ANOVA.

We confirmed the sex chromosome differences in statin regulation of representative cholesterol synthetic genes by real-time PCR. Mice with XY sex chromosomes upregulated *Hmgcs*, *Hmgcr*, *Mvk*, and *Sqle* with statin treatment compared to chow diet controls, while mice with XX sex chromosomes did not (Fig. 7B–E). *Hmgcr* expression also showed a significant gonad effect with higher expression in mice with ovaries compared to testes, irrespective of sex chromosome type and statin treatment (Fig. 7C). Our results reveal that sex differences in statin regulated gene expression depend on the individual gene, and specific genes may be influenced by gonadal sex, chromosomal sex, or a combination of the two.

## DISCUSSION

Differences between women and men have been characterized in plasma lipid profiles, hepatic lipid metabolism, and response to pharmaceutical approaches to reduce CVD risk [3,4]. There is, however, limited understanding of the molecular mechanisms that underlie these sex differences. We used *Apoe*^−/−^ FCG mice to investigate how gonadal and chromosomal sex independently impact hepatic gene expression in a hypercholesterolemic state and in response to statin. We focused on liver because of its central role in regulating homeostatic lipid levels and as the primary target for statin inhibition of cholesterol synthesis. Previous studies have identified significant sex differences in the hepatic transcriptome [26–28]. Our study extends and augments previous work by assessing the role of sex in hepatic gene regulation in physiological states that are relevant to human disease—hypercholesterolemia and statin treatment.

A general finding in our study is the demonstration that in hypercholesterolemic mice, the presence of ovaries vs. testes leads to differential expression of ~6–12% of genes on each of the 19 mouse autosomes, and presence of XX vs. XY chromosomes confers differential expression of ~3–6% of genes on each autosome. Several key findings related to the role of sex components and gene expression in hypercholesterolemic or statin-treated states also emerged. For example, an analysis of specific biological pathways that are influenced by sex components revealed that in the hypercholesterolemic state, gonadal sex influences the regulation of ER stress, whereas chromosomal sex and gonadal sex influence fatty acid metabolism. We hypothesized that the distinct sex-specific gene expression profiles in the liver of hypercholesterolemic mice would influence their response to external factors, such as statin drug treatment. Indeed, we identified strong sex-dependent responses to statin. In particular, the presence of XY chromosomes was associated with more robust upregulation of the cholesterol biosynthesis pathway genes upon statin treatment. Further analysis confirmed the absence of this statin-induced response in XX mice, irrespective of gonad type. These data demonstrate that the sex chromosome complement, independent of gonadal sex, may be an important determinant of sex-dependent statin drug response. Ultimately, the complex interplay between gonadal and chromosomal sex leads to a specific gene expression environment that is expected to determine sex differences in traits related to CVD risk, such as circulating plasma lipid levels.

Overall, our findings reveal independent effects of gonadal and chromosomal sex on the hepatic transcriptome to create sex-specific cellular phenotypes. While gonad type had the largest impact on hepatic gene expression in the conditions we assessed, we demonstrated that the sex chromosomes influence the hepatic response to statin treatment, which may have a profound impact on differential statin efficacy, effectiveness, or side effects between women and men. This may be important after middle age (when statin drugs are most commonly prescribed), as chromosomal sex effects persist after gonadal hormone levels wane. These findings join other recent data that demonstrate the integral role of sex chromosomes on cellular and whole-body physiology including mRNA and microRNA expression levels, lipoprotein metabolism, atherosclerosis, obesity, autoimmune, pulmonary, and neurological diseases [11,14,29–34]. Our study is the first to evaluate how gonadal and chromosomal sex impact cellular response to a drug treatment. An understanding of how sex components influence the response to disease conditions (such as hypercholesterolemia) and commonly prescribed drugs (such as statins) will lead to optimal treatment for all individuals.

## DECLARATIONS

### Ethics Approval and Consent to Participate

All mouse studies were conducted in accordance with UCLA Institutional Animal Research Committee (IACUC) approval.

### Consent for publication

All authors have provided consent for publication.

### Availability of data and materials

The RNA-sequencing has been deposited to Gene Expression Omnibus (GEO) database under accession <pending assignment> at http://www.ncbi.nlm.nih.gov/project/geo/.

### Competing Interests

The authors have no competing interests.

### Funding

These studies were supported by P50 GM115318 (KR), U54 DK120342 (KR), and R21 AR077782 (KR and PZ) from the National Institutes of Health, and 20POST35100000 (CBW) post-doctoral fellowship from American Heart Association.

### Author Contributions

CBW, ZWA, and KR designed the study analysis and wrote the manuscript. PZ performed mouse studies and generated RNA-seq data. CBW and ZWA analyzed transcriptome data and performed real-time PCR studies.

## Notes

### Competing Interest Statement

The authors have declared no competing interest.

## REFERENCES

1. Tsao CW, Aday AW, Almarzooq ZI, Alonso A, Beaton AZ, Bittencourt MS, et al. Heart Disease and Stroke Statistics-2022 Update: A Report From the American Heart Association. Circulation. 2022 Feb;145(8):e153–639.

2. Regitz-Zagrosek V, Kararigas G. Mechanistic pathways of sex differences in cardiovascular disease. Physiol Rev. 2017 Jan;97(1):1–37.

3. Reue K, Wiese CB. Illuminating the Mechanisms Underlying Sex Differences in Cardiovascular Disease. Circ Res. 2022;130(12):1747–62.

4. Mauvais-Jarvis F, Berthold HK, Campesi I, Carrero JJ, Dakal S, Franconi F, et al. Sex-and gender-based pharmacological response to drugs. Pharmacol Rev. 2021;73(2):730–62.

5. Goodarzi MO, Li X, Krauss RM, Rotter JI, Chen YDI. Relationship of sex to diabetes risk in statin trials. Diabetes Care. 2013 Jul;36(7):e100–1.

6. Hopewell JC, Offer A, Haynes R, Bowman L, Li J, Chen F, et al. Independent risk factors for simvastatin-related myopathy and relevance to different types of muscle symptom. Eur Heart J. 2020 Sep;41(35):3336–42.

7. Nguyen P, Leray V, Diez M, Serisier S, Le Bloc’h J, Siliart B, et al. Liver lipid metabolism. J Anim Physiol Anim Nutr (Berl). 2008 Jun;92(3):272–83.

8. Moslehi A, Hamidi-Zad Z. Role of SREBPs in Liver Diseases: A Mini-review. J Clin Transl Hepatol. 2018 Sep;6(3):332–8.

9. Istvan ES, Deisenhofer J. Structural mechanism for statin inhibition of HMG-CoA reductase. Science. 2001 May;292(5519):1160–4.

10. Mauvais-Jarvis F, Arnold AP, Reue K. A guide for the design of pre-clinical studies on sex differences in metabolism. Cell Metab. 2017 Jun;25(6):1216–30.

11. AlSiraj Y, Chen X, Thatcher SE, Temel RE, Cai L, Blalock E, et al. XX sex chromosome complement promotes atherosclerosis in mice. Nat Commun. 2019;10(1):2631.

12. Chen X, McClusky R, Chen J, Beaven SW, Tontonoz P, Arnold AP, et al. The number of X chromosomes causes sex differences in adiposity in mice. PLoS Genet. 2012;8(5):e1002709.

13. Mehrabian M, Qiao JH, Hyman R, Ruddle D, Laughton C, Lusis AJ. Influence of the apoA-II gene locus on HDL levels and fatty streak development in mice. Arterioscler Thromb a J Vasc Biol. 1993 Jan;13(1):1–10.

14. Link JC, Wiese CB, Chen X, Avetisyan R, Ronquillo E, Ma F, et al. X chromosome dosage of histone demethylase KDM5C determines sex differences in adiposity. J Clin Invest. 2020;130(11):5688–702.

15. Love MI, Huber W, Anders S. Moderated estimation of fold change and dispersion for RNA-seq data with DESeq2. Genome Biol. 2014;15(12).

16. Kuleshov M V., Jones MR, Rouillard AD, Fernandez NF, Duan Q, Wang Z, et al. Enrichr: a comprehensive gene set enrichment analysis web server 2016 update. Nucleic Acids Res. 2016;44(W1):W90–7.

17. Chen EY, Tan CM, Kou Y, Duan Q, Wang Z, Meirelles GV, et al. Enrichr: interactive and collaborative HTML5 gene list enrichment analysis tool. BMC Bioinformatics. 2013 Apr;14:128.

18. Xie Z, Bailey A, Kuleshov M V, Clarke DJB, Evangelista JE, Jenkins SL, et al. Gene Set Knowledge Discovery with Enrichr. Curr Protoc. 2021;1(3):e90.

19. Emini Veseli B, Perrotta P, De Meyer GRA, Roth L, Van der Donckt C, Martinet W, et al. Animal models of atherosclerosis. Eur J Pharmacol. 2017 Dec;816:3–13.

20. Love MI, Huber W, Anders S. Moderated estimation of fold change and dispersion for RNA-seq data with DESeq2. Genome Biol. 2014;15(12).

21. Fu S, Watkins SM, Hotamisligil GS. The role of endoplasmic reticulum in hepatic lipid homeostasis and stress signaling. Cell Metab. 2012 May;15(5):623–34.

22. Lemmer IL, Willemsen N, Hilal N, Bartelt A. A guide to understanding endoplasmic reticulum stress in metabolic disorders. Mol Metab. 2021 May;47:101169.

23. Gillespie M, Jassal B, Stephan R, Milacic M, Rothfels K, Senff-Ribeiro A, et al. The reactome pathway knowledgebase 2022. Nucleic Acids Res. 2022 Jan;50(D1):D687–92.

24. Brown MS, Goldstein JL. Multivalent feedback regulation of HMG CoA reductase, a control mechanism coordinating isoprenoid synthesis and cell growth. J Lipid Res. 1980 Jul;21(5):505–17.

25. Jiang SY, Li H, Tang JJ, Wang J, Luo J, Liu B, et al. Discovery of a potent HMG-CoA reductase degrader that eliminates statin-induced reductase accumulation and lowers cholesterol. Nat Commun. 2018 Dec;9(1):5138.

26. Yang X, Schadt EE, Wang S, Wang H, Arnold AP, Ingram-Drake L, et al. Tissue-specific expression and regulation of sexually dimorphic genes in mice. Genome Res. 2006 Aug;16(8):995–1004.

27. Blencowe M, Chen X, Zhao Y, Itoh Y, McQuillen CN, Han Y, et al. Relative contributions of sex hormones, sex chromosomes, and gonads to sex differences in tissue gene regulation. Genome Res. 2022 May;32(5):807–24.

28. Goldfarb CN, Karri K, Pyatkov M, Waxman DJ. Interplay between GH-regulated, sex-biased liver transcriptome and hepatic zonation revealed by single nucleus RNAseq. Endocrinology. 2022 Jul;163(7):bqac059.

29. Arnold AP. Four Core Genotypes and XY* mouse models: Update on impact on SABV research. Neurosci Biobehav Rev. 2020 Dec;119:1–8.

30. Link JC, Hasin-Brumshtein Y, Cantor RM, Chen X, Arnold AP, Lusis AJ, et al. Diet, gonadal sex, and sex chromosome complement influence white adipose tissue miRNA expression. BMC Genomics. 2017 Dec 17;18(1):89.

31. Link JC, Chen X, Prien C, Borja MS, Hammerson B, Oda MN, et al. Increased high-density lipoprotein cholesterol levels in mice with XX versus XY sex chromosomes. Arterioscler Thromb Vasc Biol. 2015 Aug;35(8):1778–86.

32. Itoh Y, Golden LC, Itoh N, Matsukawa MA, Ren E, Tse V, et al. The X-linked histone demethylase Kdm6a in CD4+ T lymphocytes modulates autoimmunity. J Clin Invest. 2019;129(9):3852–63.

33. Davis EJ, Broestl L, Williams G, Garay BI, Lobach I, Devidze N, et al. A second X chromosome contributes to resilience in a mouse model of Alzheimer’s disease. Sci Transl Med. 2020;12(558):eaaz5677.

34. Cunningham CM, Li M, Ruffenach G, Doshi M, Aryan L, Hong J, et al. Y-Chromosome Gene, Uty, Protects Against Pulmonary Hypertension by Reducing Proinflammatory Chemokines. Am J Respir Crit Care Med. 2022 Jul;206(2):186–96.

